# A new clustering approach identifies tumor-specific common TCRs with pan-cancer reactivity

**DOI:** 10.1101/2025.04.26.650660

**Authors:** S. Hennig, K. Gennermann, S. Elezkurtaj, V. Seitz, B. Hirsch, A. Droege, S. Schaper, D. Bents, S. Eggeling, C. Beushausen, H. Herbst, N. Genzel, J. Glökler, C. Wölfel, C. Doppler, V. Lennerz, R. Hammer

## Abstract

Tumor-specific T-cells are key in combating cancer as shown in adoptive cell therapy with tumor infiltrating lymphocytes (TILs) and checkpoint inhibitor therapy. Studies in many types of cancer have shown that preexisting tumor reactive T-cells are not only tumor-but typically also patient-specific, requiring personalized treatment options. For viral infections, public T-cell receptors (TCRs) with substantial sequence homologies suggest shared immune-dominant targets in human leucocyte antigen (HLA)-matched individuals. We hypothesized that also in the complex TCR repertoires of tumors subsets of tumor-specific TCRs exist that can be found in different patients with identical or near identical TCRs. This paper presents a TCR-V(D)J-sequence clustering approach identifying clusters of tumor-specific common TCRs mainly from TILs of non-small cell lung cancer (NSCLC) patients. Using two TCR-clusters as examples, we show that T-cells engineered genetically to only express those common TCRs recognized HLA-matched allogeneic tumor cell lines in a cluster typical manner. Recognition of allogeneic tumors was dependent on the HLA allele inferred by the cluster and could be blocked by HLA antibodies. In addition to NSCLC, TCR repertoire analyses in pancreatic ductal adenocarcinoma and a smaller number of breast and colorectal cancer samples revealed TCRs highly homologous or even identical to NSCLC cluster TCRs. TCR-T cells expressing TCRs from these tumors assigned to a specific cluster recognized allogeneic tumor lines in the expected cluster-typical manner. These findings suggest a pan-cancer therapeutic potential of tumor-specific common TCRs.

## Introduction

In contrast to the chimeric antigen receptor (CAR-T) approach, T-cell receptor (TCR)-T therapy addresses intracellular tumor antigens thereby holding great promise for the treatment of a broader range of cancers, including solid tumors (Klebanoff 2023, Peri 2023)[1, 2]. During recent years, antigen-agnostic methods for the prediction of tumor-reactive T-cells have been proposed (Lowery 2022, Meng 2023, Tan 2024, Pétremand 2024, Lennerz 2025)[3-6]. Such approaches have described the identification of tumor-reactive T-cells in individual patients as the starting point for novel personalized TCR-T therapies. As the development of individual treatments for each single patient imposes formidable challenges, we asked whether identical or near identical tumor-specific TCRs may exist that can be found in a significant proportion of cancer patients. To search for such common TCRs we have developed a novel TCR-V(D)J-sequence clustering pipeline.

In this paper we describe a new clustering approach which identifies tumor-specific common TCRs mainly in NSCLC patients. Tumor specificity of T-cell clonotypes is determined per patient by comparative TCR-repertoire sequencing of TILs and T-cells from adjacent normal tissue and indicated by significant enrichment of TCRs in the tumor (Lennerz 2025)[6]. Common tumor-specific TCRs are identified by a novel clustering method employing TCR-V(D)J-sequences of paired chains (α/β) of the T-cell receptor.

## Results

### Clustering of tumor-specific paired TCRs

Employing a novel pipeline of clustering algorithms (for details see Materials and Methods), we used TCR-repertoire profiling (TCRSeq) data from a database of >1.7 Mio TCR sequences from tumor-infiltrating T-cells (TILs) and from T-cells found in adjacent normal tissues for the identification of tumor-specific TCRs being highly homologous between different patients and even distinct cancer types. The clustering results in this paper are based on the data source described in Table 1.

**Table 1:** Details of patient cohorts with respect to paired (α/β) TCR sequence data from tumors. Both chains of TCRs were actively used for TCR clustering.

|  | Lung | Pancreas | Breast | Colon |
| --- | --- | --- | --- | --- |
| Number of Patients | 62 | 10 | 5 | 7 |
| Number of paired TCRs (tumor) | 171929 | 12582 | 8037 | 10947 |

Our approach is - to the best of our knowledge – unique in that it builds clusters of TCRs with a priori evidence of their tumor-specificity and with fully resolved sequences of paired (α/β) TCR chains. It is completely antigen-agnostic, i.e. no assumptions about common antigens are made or necessary. In addition, while there are several published TCR clustering algorithms (Glanville 2017, Bo Li 2018, Mayer-Blackwell 2021, Zhang 2021)[7-10], only few will make use of both chains of the TCR simultaneously (Wang 2022)[11] and none is using tumor-specificity so far.

Figure 1 shows a schema of our clustering approach which is described in detail below (Materials and Methods). Core of the procedure is the use of well characterized TCR-sequences identified by a combination of bulk CD8 TCR sequencing and single-cell VDJ sequencing (10X genomics) of TILs isolated from fresh NSCLC surgery samples. Our general method for the antigen-agnostic identification of tumor-specific TCRs has recently been published (Lennerz, 2025)[6].

**Figure 1:**
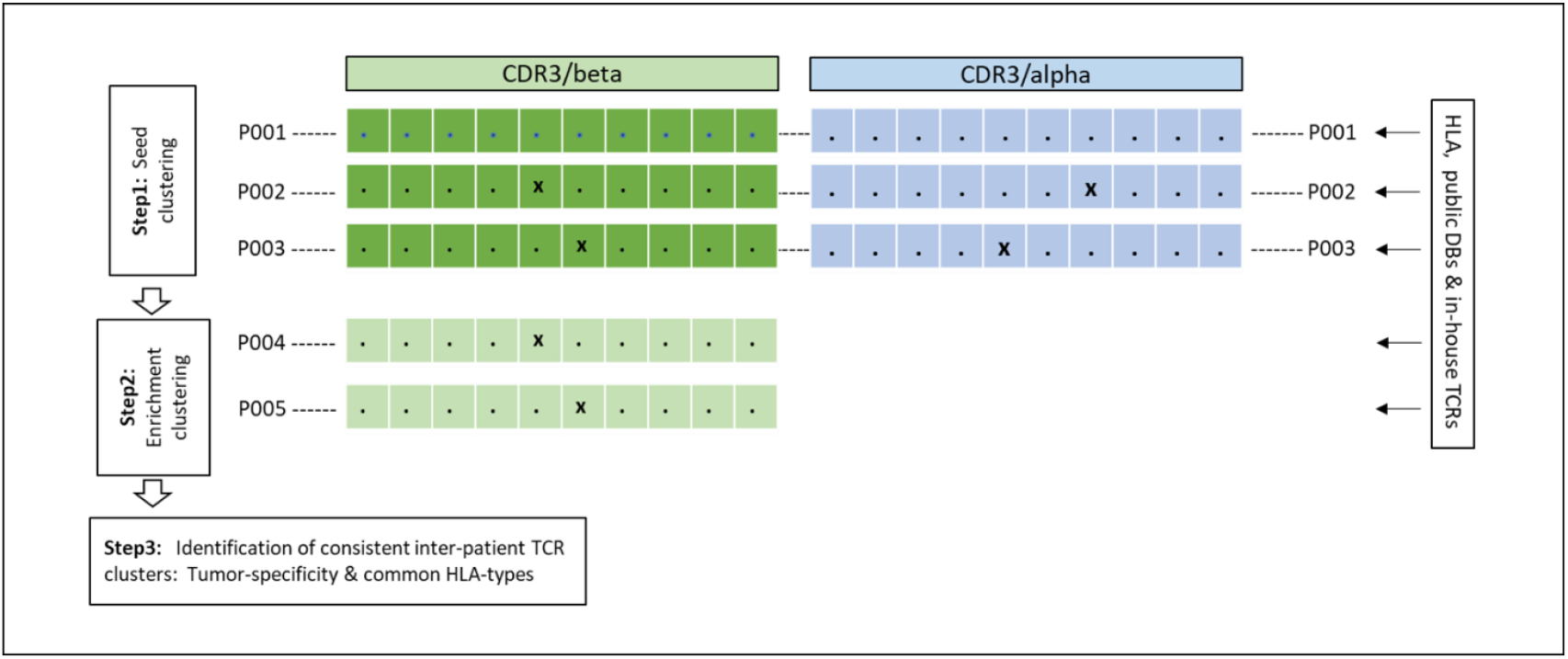
Schema of tumor-specific inter-patient TCR clustering. **Step1:** A fast sequence comparison method is used to find identical or very similar TCRs between distinct patients. We use a distance metrics like the Levenshtein distance counting the number of mismatches between combined alpha and beta chains (see below). The dots denote identical Aa, the, x’ are symbolizing Aa differing from the consensus. **Step2:** Once the alpha/beta-paired sequences of all TCRs from patients’ TILs are grouped in seed clusters the clusters are enriched by available TCR-beta sequences with no knowledge about alpha-chains. **Step3:** In the final step the sequence based clusters are re-analyzed with respect to HLA enrichment and tumor-specificity of cluster TCRs.

### Examples of αβ-TCR clusters from tumors of NSCLC patients with tumor-specific enrichment

Our clustering method enabled the identification of TCR clusters with tumor-specific enrichment. Table 2 and Table 3 depict Cluster A and B as examples for tumor-specific clusters. The alpha and beta chains of all paired TCRs found in our patient cohort for these two clusters are shown.

**Table 2:**
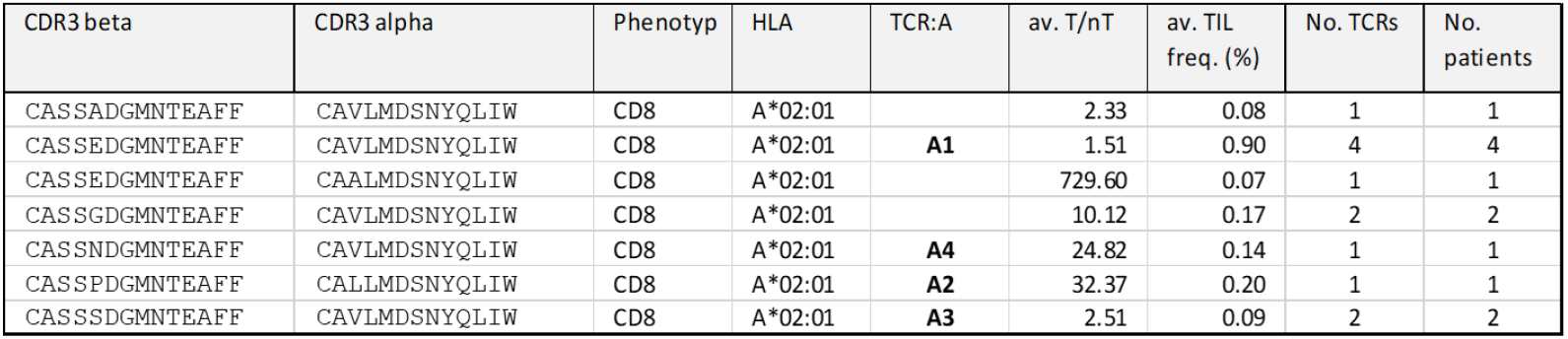
Cluster A contains 7 distinct TCRs from 10 different NSCLC patients. TCRs identical between patients or within a patient are shown with their average frequencies and tumor vs non-tumor (T/nT) ratios. Median values of T/nT ratios and TIL frequencies (%) are = 10.1 and 0.14%, respectively. The clonotypes A1-A4 were subjected to extensive functional validations (see below). All 10 patients were found to share the HLA-A*02:01 allele.

| CDR3 beta | CDR3 alpha | Phenotyp | HLA | TCR:A | av. T/nT | av. TIL freq. (%) | No. TCRs | No. patients |
| --- | --- | --- | --- | --- | --- | --- | --- | --- |
| CASSADGMNTEAFF | CAVLMDSNYQLIW | CD8 | A*02:01 |  | 2.33 | 0.08 | 1 | 1 |
| CASSEDGMNTEAFF | CAVLMDSNYQLIW | CD8 | A*02:01 | **A1** | 1.51 | 0.90 | 4 | 4 |
| CASSEDGMNTEAFF | CAALMDSNYQLIW | CD8 | A*02:01 |  | 729.60 | 0.07 | 1 | 1 |
| CASSGDGMNTEAFF | CAVLMDSNYQLIW | CD8 | A*02:01 |  | 10.12 | 0.17 | 2 | 2 |
| CASSNDGMNTEAFF | CAVLMDSNYQLIW | CD8 | A*02:01 | **A4** | 24.82 | 0.14 | 1 | 1 |
| CASSPDGMNTEAFF | CALLMDSNYQLIW | CD8 | A*02:01 | **A2** | 32.37 | 0.20 | 1 | 1 |
| CASSSDGMNTEAFF | CAVLMDSNYQLIW | CD8 | A*02:01 | **A3** | 2.51 | 0.09 | 2 | 2 |

**Table 3:** Cluster B contains 8 distinct TCRs from 9 different NSCLC patients. TCRs identical between patients or within a patient are shown with their average frequencies and T/nT ratios. Median values of T/nT ratios and TIL frequencies (%) are 5.2 and 0.27%, respectively. The clonotypes B1-B4 were subjected to extensive functional validations (see below). All 9 patients were found to share the HLA-B*08:01 allele.

| CDR3 beta | CDR3 alpha | Phenotyp | HLA | TCR:B | av. T/nT | av. TIL freq. (%) | No. TCRs | No. patients |
| --- | --- | --- | --- | --- | --- | --- | --- | --- |
| CASSAGPNYEQYF | CAASFNQGGKLIF | CD8 | B*08:01 | **B1** | 1709.30 | 0.17 | 1 | 1 |
| CASSQGPNYEQYF | CAASINQGGKLIF | CD8 | B*08:01 |  | 1.43 | 0.16 | 1 | 1 |
| CASSLGPNYEQYV | CAASSNQGGKLIF | CD8 | B*08:01 |  | 2.10 | 1.52 | 1 | 1 |
| CASSPGPNYEQYF | CAGAYNQGGKLIF | CD8 | B*08:01 | **B2** | 205.30 | 0.02 | 1 | 1 |
| CASSPGPNYEQYF | CAASINQGGKLIF | CD8 | B*08:01 | **B4** | 1.269 | 0.644 | 4 | 2 |
| CASSPGPNYEQYF | CAASYNQGGKLIF | CD8 | B*08:01 | **B3** | 10.22 | 1.00 | 3 | 2 |
| CASSPGPNYEQYF | CAASWNQGGKLIF | CD8 | B*08:01 |  | 8.27 | 0.24 | 4 | 1 |
| CASSQGPNYEQYF | CAASYNQGGKLIF | CD8 | B*08:01 |  | 1.70 | 0.30 | 3 | 3 |

It is evident from these two examples that our clustering algorithm has collected highly homologous clusters of paired TCR sequences with constant lengths and only minimal changes of alpha- and beta-chains. Although the ratio between tumor and non-tumor (T/nT) is variable, it is obvious that the two clusters are showing tumor-specific enrichment (Tables 2 and 3) with median enrichment factors of 10.1 and 5.2 for Cluster A and B, respectively. A striking output of the TCR sequence-based clustering is the strong association of both clusters with specific HLA alleles (Tables 2 and 3). HLAs are not part of the clustering method. On the contrary, the HLA-associations are obtained as results of the clustering. Cluster A and B show a highly significant HLA-association with HLA-A*02:01 (p-val. 0.001) and HLA-B*08:01 (p-val. 0.0003), respectively.

### TCRs of tumor-specific Clusters A and B recognize HLA-matched allogeneic tumor lines

The two tumor-specific TCR-clusters A and B depicted in Tables 2 and 3 are characterized by well-defined HLA-restrictions and paired TCR-sequences with almost identical CDR3-regions found in TILs of different NSCLC patients. For this reason, we hypothesized, that the cluster TCRs share specificity for common antigens and therefore might have the ability to recognize allogeneic HLA-matched tumors. We selected four TCRs (A1-A4, B1-B4) from each cluster to be used for response analyses against allogeneic tumor cell lines (see Tables 2 and 3). The TCRs were transduced into healthy donor T-cells by retroviral transduction. To eliminate potential alloreactivity of transduced donor T-cells against target cell lines, T-cells were depleted from their endogenous TCRs by CRISPR/CAS9-mediated knockout (KO) as described in (Lennerz, 2025)[6].

Figure 2 shows an ELISpot assay testing TCR-T cells transduced with cluster A TCRs A1-A4 against five HLA-A*02:01-positive NSCLC cell lines. All four TCRs showed reactivity against the cell lines NCI-H1703, NCI-H1792 and MZ-LC-16, demonstrating their ability to recognize allogeneic tumors. On the other hand, they failed to respond against MOR/CPR and NCI-H661, with the exception of TCR A1 that showed a modest response against MOR/CPR. Further testing was carried out with chronic myeloid leukemia (CML) cell line K562. The Cluster A TCRs induced responses when K562 cells were transduced with HLA-A*02:01 but revealed no recognition when tested against untransduced K562 cells (not shown). Reactivity of Cluster A TCRs against MZ-LC-16 was blocked by an anti-HLA-A*02 antibody indicating a peptide-HLA restricted mechanism (Figure 3).

**Figure 2:**
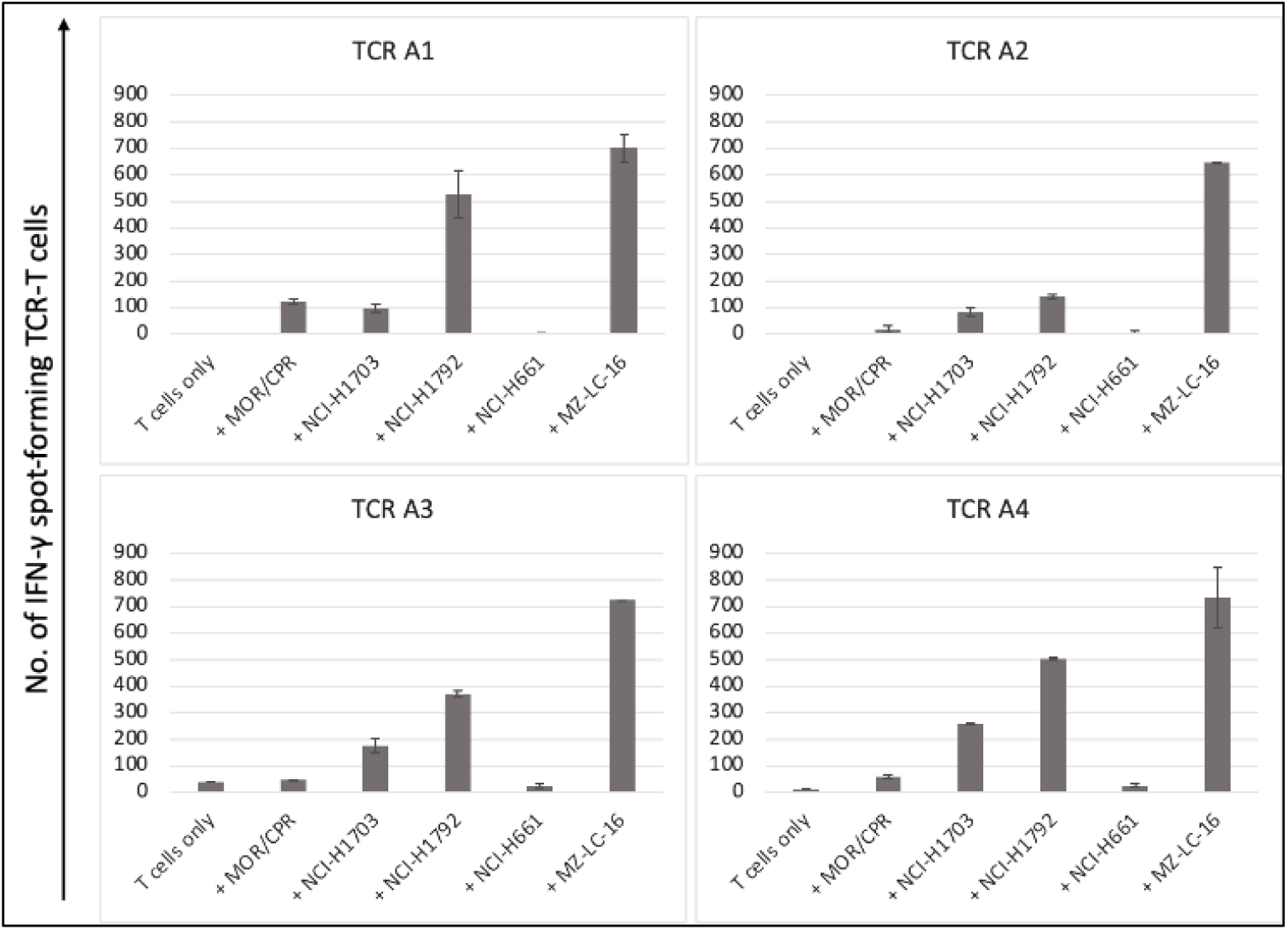
CD8+ T-cells from a healthy donor were depleted from endogenous TCRs by CRISPR/CAS9-mediated knockout and transduced with Cluster A TCRs (A1 - A4). Transduced TCR-T cells were tested against five HLA-A*02:01 positive NSCLC cell lines (MOR/CPR, NCI-H1703, NCI-H1792, NCI-H661 and MZ-LC-16) by IFN-γ-Elispot assay. All reactions were performed in duplicates, mean values are shown, error bars show standard deviations.

**Figure 3:**
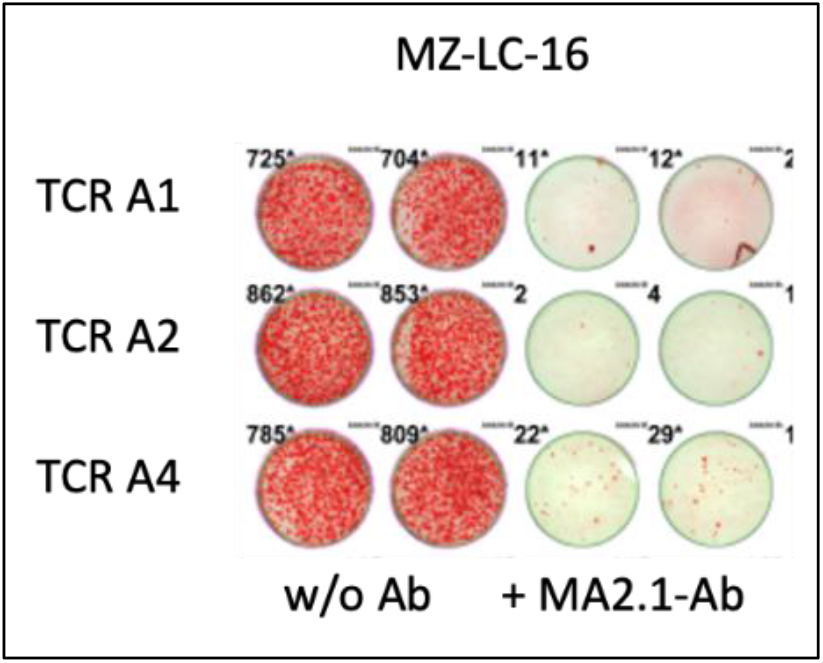
IFN-γ-Elispot assay of TCR-T cells transduced with TCR A1, TCR A2 and TCR A4 against NSCLC cell line MZ-LC-16 with and without anti-HLA-A*02:01 antibody MA2.1. All reactions were performed in duplicates. Spot numbers are shown on top of the wells.

Figure 4 depicts the screening results of TCR B3 transduced TCR-T cells against seven NSCLC cell lines. TCR B3 clearly recognized NCI-H1703, this was one out of three HLA-B*08:01 carrying tumor lines tested. It is also evident that none of the four HLA-B*08:01 negative NSCLC cell lines was able to elicit a significant response. In Figure 5 all four transduced TCR-T cells of Cluster B were tested against NCI-H1703. All four TCRs of Cluster B exhibited robust recognition. In experiments with K562 cells, TCR B3 was shown to recognize K562 cells transduced with HLA-B*08:01 but not untransduced K562 cells. Reactivity against B*08:01 transduced K562 cells was blocked by a pan-HLA class I antibody indicating HLA-restricted reactivity (not shown).

**Figure 4:**
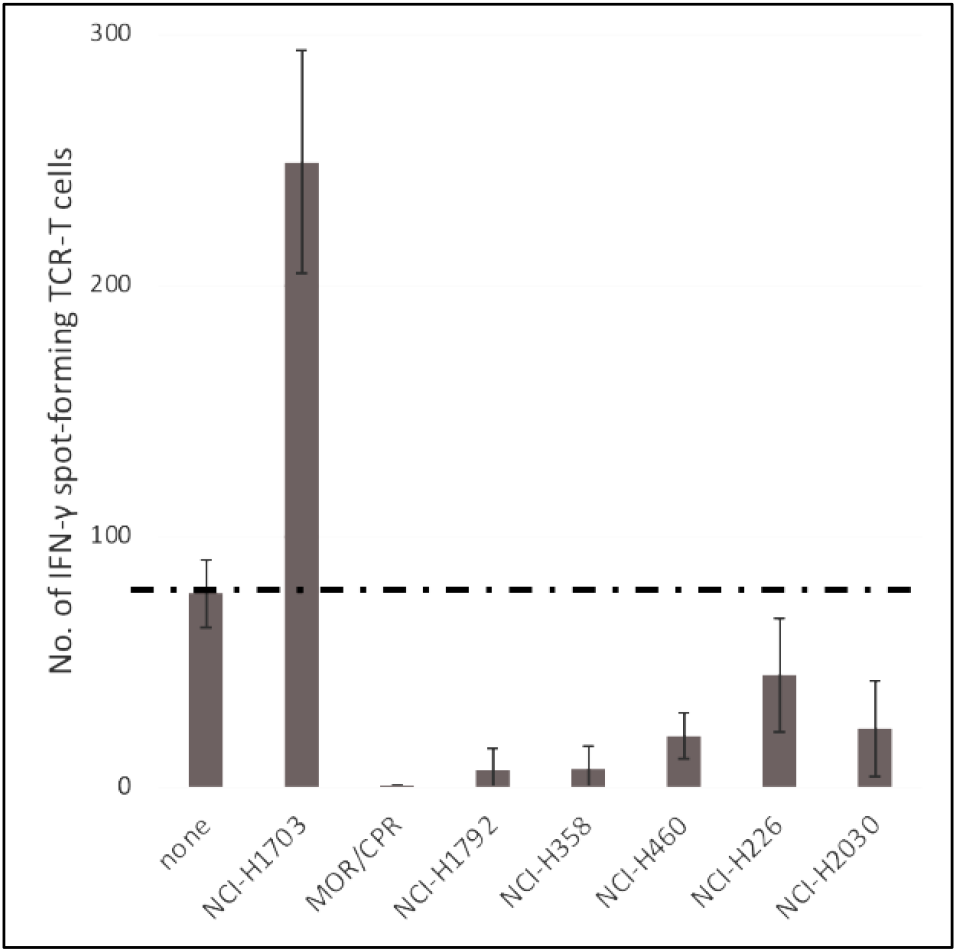
TCR-T cells transduced with TCR B3 of Cluster B were tested against seven NSCLC cell lines. NCI-H1703, MOR/CPR and NCI-H1792 express the cluster-relevant HLA-B*08:01 allele. NCI-H358, NCI-H460, NCI-H226 and NCI-H2030 do not carry the HLA-B*08:01 allele. All ELISpot reactions were set up in duplicates, mean values are shown, error bars show standard deviations.

**Figure 5:**
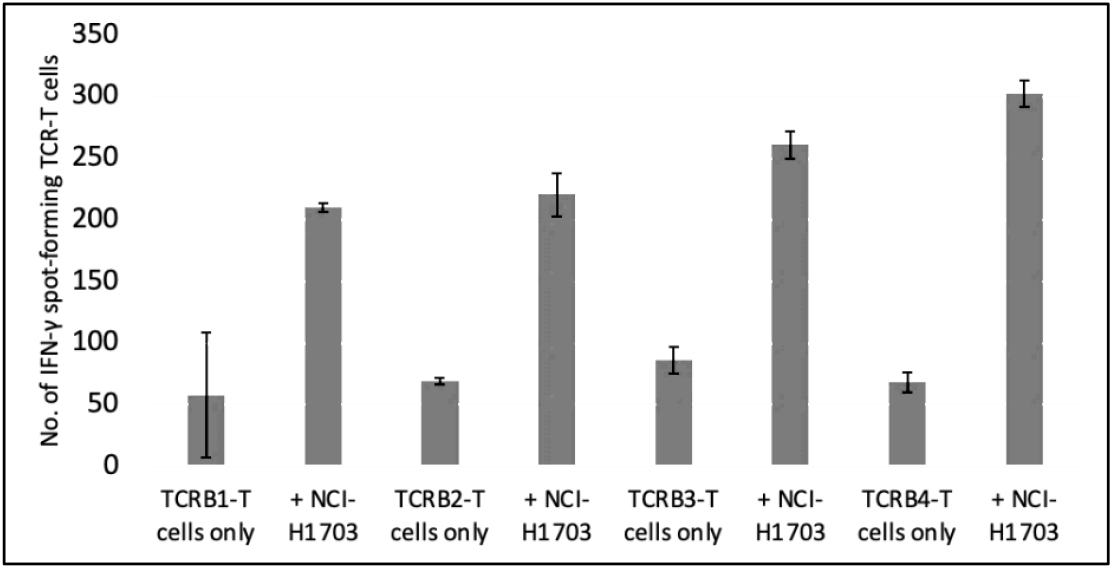
CD8+ T-cells from a healthy donor were depleted from endogenous TCRs by CRISPR/CAS9-mediated knockout and transduced with Cluster B TCRs (B1 - B4). Transduced TCR-T cells were tested against HLA-B*08:01 positive NSCLC cell line NCI-H1703. IFN-γ-Elispot reactions were performed in duplicates, mean values are shown, error bars show standard deviations.

In summary, our results with TCR-T cells expressing tumor-specific cluster TCRs derived from NSCLC patients show that they were able to recognize HLA-matched allogeneic tumor lines. TCRs of a given cluster recognized the same subset of allogeneic tumor cells. Recognition was dependent on the HLA allele defined by the TCR cluster and blocked by anti-HLA antibodies.

### Further screening for cluster TCRs in tumors of breast, colorectal and pancreatic cancer

We set out to explore whether it is feasible to find common cluster TCRs in tumors other than NSCLC. Table 1 summarizes the investigated tumor-types, patient numbers and input data for clustering.

Table 4 shows that we identified TCRs assignable to clusters A and B in breast, colorectal and pancreatic cancer. In breast cancer we found TCR Ma25, a receptor identical in its CDR3 amino acid sequences to TCR A1 of Cluster A. The tumor cell reactivity of TCR A1 has been documented in Figure 2. In colorectal cancer we identified CRC18 (Table 4). This TCR is a perfect fit for Cluster B (Table 3). CRC18 has only one amino acid change in CDR3beta and an identical CDR3alpha in comparison to other members of Cluster B.

**Table 4:**
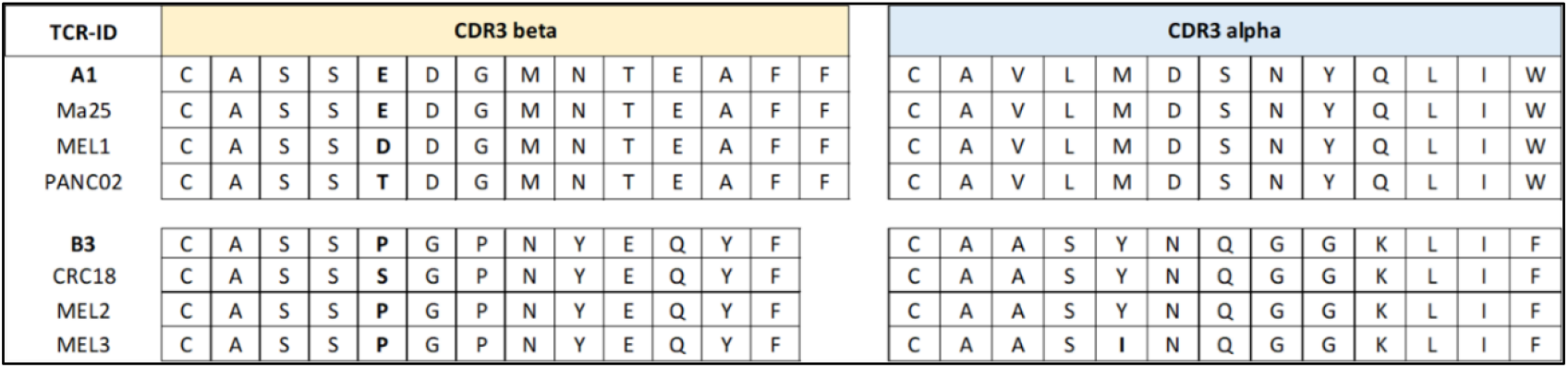
Pan-cancer clusters A and B of paired TCRs derived from tumors of lung, breast, colon, pancreas and melanoma. Variable amino acids of CDR3beta and CDR3alpha are highlighted. TCRs in Cluster A and B are restricted by HLA-A*02:01 or HLA-B*08:01, respectively.

In pancreatic cancer we identified PANC02 (Table 4). Thus far, this receptor that is highly homologous to Cluster A TCRs (Table 2) has not been detected by us in NSCLC patients. In Figure 6 functional tests with PANC02 are reported. When tested against five NSCLC cell lines PANC02 exhibited a comparable response profile to that observed with all other cluster A TCRs (compare Figure 6 with Figure 2). Thus, a TCR identified and cloned from TILs of a pancreatic cancer patient effectively recognized different NSCLC-cancers suggesting pan-cancer reactivity.

**Figure 6:**
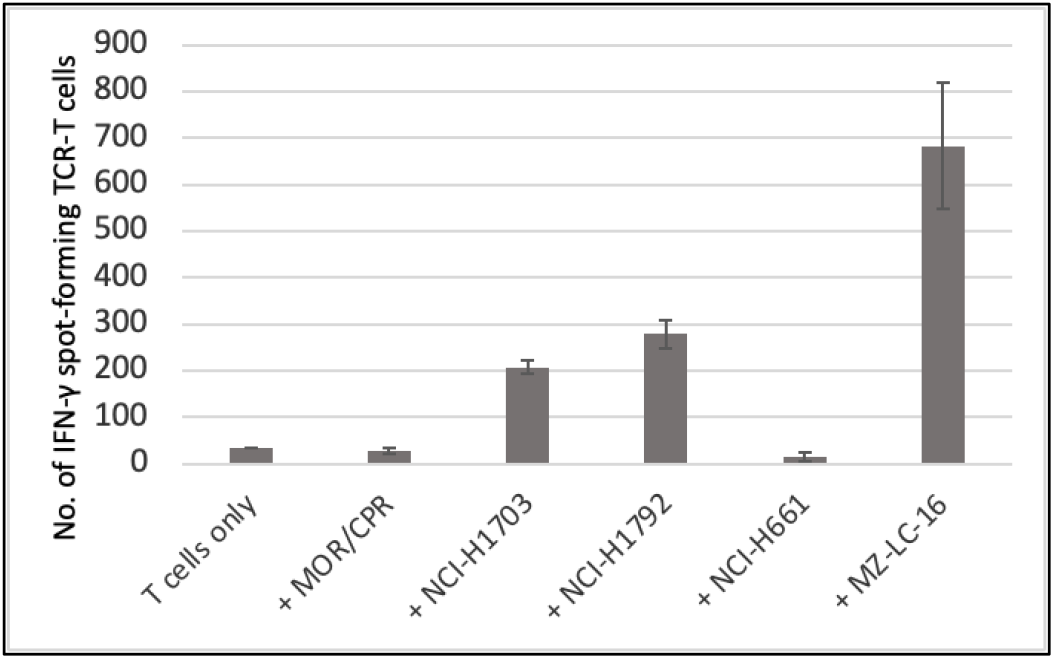
TCR-T cells transduced with the pancreatic TCR PANC02 were tested against five HLA-A*02:01 positive NSCLC cell lines (MOR/CPR, NCI-H1703, NCI-H1792, NCI-H661 and MZ-LC-16) by IFN-γ-Elispot assay. The results correspond to the recognition profile of other cluster A TCRs. All reactions were set up in duplicates, mean values are shown, error bars show standard deviations.

Complementary to our own TCR-repertoire sequencing in different cancers we became aware of two recent publications on cancer related TCR repertoires in cohorts of melanoma and pediatric brain tumors. In an informative study in melanoma tumor-derived T-cells of 32 patients treated with checkpoint inhibitors had been subjected to single-cell sequencing analysis (Sade-Feldman 2018)[12]. Among the reported paired CD8 TCRs we have identified clear hits for both Cluster A and Cluster B (Table 4). MEL 1 is an almost identical clonotype to Cluster A TCRs with one amino acid differing only. MEL 2 and MEL 3 are identical in their amino acid sequences to TCR B3 and TCR B4 of Cluster B. Both receptors have been functionally tested. Both TCRs do recognize HLA-matched allogeneic tumor cells as shown in Figure 5.

In a very comprehensive study on 996 patient samples of pediatric brain tumors (Raphael and Kohanbash 2024)[13] the authors identified groups (clusters) of TCRs with significant tumor specificities and sequence homologies between different individuals. Although they give only TCR-beta sequences of their clusters we found an astonishing number of them being closely homologous to our cluster TCRs. These are very recent findings which are currently under further investigation (no data shown here).

In summary, our results show that common TCRs identified in NSCLC patients can also be detected in other solid tumors, like breast, colorectal, pancreatic, skin and even pediatric brain cancers. Thus, we have *de facto* extended inter-patient TCR clustering to inter-cancer TCR clustering. This is in line with the functional capability of tumor-specific common TCRs to recognize allogeneic tumors in HLA-matched settings. Our findings suggest an exciting pan-cancer therapeutic potential of tumor-specific common TCRs.

## Discussion

We hypothesized that besides the well-established antitumor T-cell immunity arising in tumor-draining lymph nodes and delineated in the classical cancer-immune cycle model (Chen, Mellman 2013)[14], a significant part of the anticancer T-cells found in tumor probes have not originated in lymph nodes. In contrast, they were generated within the tumor microenvironment (TME) originating from naïve T-cells or resident pathogen-specific memory T-cells that encountered their cognate tumor antigens followed by proliferation and tumor-specific enrichment of these clones (Gueguen 2021)[15].

In fact, there is scientific support for antitumor T-cell immunity generated in the TME. Since the seminal paper in 2008 (Dieu-Nosjean 2008)[16], several studies have described the existence of postnatally induced tertiary lymphoid structures (TLSs) in the tumor and in the cancer invasive area resembling lymph nodes without encapsulation and containing B-cells, T-cells and dendritic cells (Goc 2014, Hiraoka 2016, Schumacher 2022)[17-19]. Whilst some reports were controversial (Kang 2021)[20], most papers evaluate TLSs as drivers of the antitumor response (Dieu-Nosjean 2016)[21]. In fact, proliferation marker Ki-67 positive cells within TLSs are observed, indicative of ongoing cell division (Poschke 2016)[22]. There is even the reasoning of a potentially faster and more efficient immune response of TLSs in comparison to distant lymph nodes due to the close proximity of immune and cancer cells (Schumacher 2022)[19].

We set out to search for T-cell clones enriched in the tumor. To trace such clones we used TCR repertoire immune-sequencing methods at single clonotype resolution (Seitz 2015)[23]. Specifically, we interpreted the preferred tumor localization of an individual clone as determined quantitatively by the ratio of the clonotype frequencies between tumor and adjacent non-tumor tissue as evidence for its tumor specificity.

Sampling of T-cells for the identification of tumor-specific clonotypes from tumor probes is not trivial. Studies with more than 20 tumor types employing several methods for purity assessment have shown that tumor samples usually contain significant amounts of non-cancerous cells and tissues (Aran 2015)[24]. Besides fibroblasts and infiltrating immune cells, tumor cells due to their invasive nature (Carter 2012)[25] penetrate also normal (healthy) tissue. Such normal tissue, especially when derived from barrier organs, like skin, lung and intestine, is a rich source of T-cells many of them of pathogen-specific nature residing as tissue resident memory T-cells (Schenkel 2014, Purwar 2011)[26, 27]. It is well established that pathogen-specific T-cells can exhibit tumor cross-reactivity and as such can constitute important precursors of tumor-reactive T-cells (Bessel 2020, Balachandran 2017)[28, 29]. Moreover, it is postulated that tumor antigen loaded antigen presenting cells (APCs) in the tumor microenvironment can activate cross-reactive pathogen-specific memory T-cells (Lang 2022)[30] and that memory T-cells have a 50-fold lower functional activation threshold and respond more quickly than naïve T-cells (Pihlgren 1996, Lang, 2022)[30, 31]. In fact, when comparing our clustering results with the VDJdb data base (Shugay 2018)[32] it is evident that Cluster A (Table 1) contains TCRs associated with pathogen reactivity. With calculation of Cluster A by our α/β-TCR clustering approach we have confirmed findings of Chiou et al. (2021)[33] who used the GLIPH2 algorithm to cluster only TCR-β-chains of TILs. This group derived one paired cluster TCR from the tumor and showed EBV and E. coli cross reactivity and, by yeast library display, identified an overexpressed protein as the potential target antigen on tumor cells (Chiou 2021)[33].

Another issue to deserve attention are autoreactive T-cells. Recently, significant fractions of tumor reactive TCRs isolated by single cell sequencing from different murine pancreatic cancer models were shown to be autoreactive (Zens 2023, CIMT poster 297)[34]. There is also ample clinical evidence of autoimmune reactions caused by checkpoint inhibitors, which commonly impact dermatologic, gastrointestinal and endocrine systems and less frequent but often more severely affect pulmonary, neurologic and cardiac systems (Fazer-Posorske 2024)[35]. Such autoimmune side effects are mostly triggered by unblocking autoreactive T-cells present in these tissues. Autoantigens are “self-antigens”. They are part of the natural repertoire of cells and recognized by the immune system by mistake. They are not tumor specific and therefore autoreactive TCRs should be filtered out by employing the T/nT ratio. This is an inherent feature of our selection method.

In the last decade, it has been claimed that tumor-reactive T-cells from TILs can be identified simply by cell surface markers (Gros 2014, Duhen 2018)[36, 37]. More recently, the advent of advanced single cell TCR and gene expression sequencing enabled the exploration of the transcriptomic characteristics of intratumorally neoantigen-specific T-cells (Hanada 2022, Lowery 2022, Zheng 2022)[3, 38, 39] and T-cells reactive against autologous tumor lines (Oliveira 2021, Meng 2023, Tan 2024, Petremand 2024)[4, 5, 40, 41]. These studies have led to the delineation of informative gene expression signatures for tumor reactivity by different working groups, however with major differences in key genes, like TIGIT, PDCD1, TOX, LAG3 and CTLA4 between the different groups (Petremand 2024)[41].

Although being tumor-antigen-agnostic as well, our approach to trace tumor-specific T-cells is quite different since it is based on the identification of tumor-enriched clonotypes. We believe that the prediction of therapeutically relevant TCRs in terms of efficacy and safety is right now a burning issue and that combining different measures, i.e. high abundance, high tumor-to-non-tumor frequency ratio and informative gene expression signature, will be beneficial to bring more accuracy and safety into the antigen-agnostic selection of clinically relevant T-cell receptors towards the development of personalized T-cell therapies.

Whilst studies with our method in single patients allow the identification of “private” tumor-specific T-cell clonotypes as potential starting point for the development of personalized therapies by employing their TCRs for TCR-T approaches (Lennerz 2025)[6], we extended our studies to look for tumor-specific T-cell clonotypes with identical or near identical TCRs across patients in a cohort of 62 NSCLC patients employing a new clustering algorithm that incorporated the tumor-to-non-tumor clonotype ratio as the only parameter to indicate tumor-specificity.

In a previous publication (Lennerz 2025)[6] we showed that T-cell clonotypes selected by our method express several of the known and putative genetic activation markers. Especially, we could demonstrate that tumor-specific T-cells selected by our approach followed the expected differentiation trajectories consistent with cytotoxicity, chronic stimulation and exhaustion.

In the present paper we report the existence of tumor-specific TCR clusters across a significant proportion of cancer patients. These clusters are characterized by consensus TCRs with highly homologous alpha- and beta-chains in different patients who are sharing the cluster-typical HLA allele. We found that many of the cluster TCRs were not derived from the abundant tumor-specific T-cell clonotypes in the tumor. It required a sensitive method to reliably detect these tumor-specific clusters. We achieved this by combining the 10X single cell approach with our TCRsafe™ sequencing technology that uses genomic DNA extracted from CD8+ sorted T-cells and employing a threshold for detection for CD8+ TCRs of 0,001%. Clonotypes comprising less than 0.001% of all reads were excluded from the analysis due to their presumed representation of either background noise or genuine signals that were too low to be statistically compared.

Of potentially high significance is the finding that the tumor-specific common TCRs described in this study are not only relevant in the originating solid tumor type. Their relevance appears to extend to various other solid tumors. This report shows that examples of NSCLC derived tumor-specific common TCRs with proven capability to recognize HLA-matched allogeneic tumors can be detected also in surgical samples of pancreatic-, breast-, colorectal- and skin-cancer patients. Such findings encourage further studies to explore the therapeutic potential of tumor-specific common TCRs.

## Materials and Methods

### Clustering of alpha/beta paired TCR sequences from distinct patient tumors

To perform sequence comparisons and clustering of hundreds of thousands of TCRs within a short time period we are using fast basic algorithms like CD-HIT (Fu 2023) and BLASTP (Altschul 1990). E.g., CD-HIT is able to perform a high-quality clustering of around 250 000 paired TCRs within 30 seconds on a medium performance Linux PC (32 GiB RAM memory and a 16 cores AMD® Ryzen 7 4800h with radeon graphics processor).

The following basic constraints and conditions for clustering of TCR (CDR3) peptides are applied:

- Alpha- and beta-chain CDR3 peptide sequences are treated as 1 sequence. They are virtually glued together after clipping of the 2 amino acids each on the V- and J-segments. Thus, only the core variable parts of the CDR3s are used.
- These ‘glued’ paired sequences are clustered with 89% sequence similarity using CD-HIT, which results in the seed clusters.
- A consensus sequence for the beta-chain and alpha chain is derived for each cluster independently.
- A Levenshtein-distance of 2 is allowed to add further tumor-specific TCR beta or alpha repertoire data towards the beta or alpha consensus.

Fisher’s exact test was used to determine significantly enriched HLA-types for each cluster. For a meaningful determination, we only considered clusters with a minimum of 5 patients and adjusted the p-values with the Benjamini-Hochberg procedure for multiple testing.

### Patient material

From 62 patients with NSCLC, fresh tissue specimens from tumor- and adjacent normal-lung tissue were selected by pathologist and provided to the laboratory for preparation of lymphocyte subpopulations to be used for TCR-repertoire sequencing and TIL single-cell RNA-sequencing (see below). From 10 patients with pancreatic adenocarcinoma, 5 with breast cancer and 7 with colorectal cancer, fresh tumor tissue was selected and provided by the pathologist for single cell preparation used for TIL single-cell RNA-sequencing (see below). The study was performed in accordance with the declaration of Helsinki. Sample collection from patients was approved by the Ethics Committees of the Berlin Medical Association and Charité University Hospital Berlin (Eth-08/18, EA1/270/23) and signed informed consent was obtained from all patients.

### Tumor and non-tumor tissue processing

Fresh tumor and adjacent normal tissue samples from NSCLC patients were sliced using scalpels and subjected to GentleMACS tissue dissociation according to the manufacturer’s instructions (Miltenyi Biotec, Bergisch-Gladbach, Germany). Filtered through 70 μm-cell strainers, one aliquot each of the cell suspensions underwent Percoll gradient centrifugation and the remainder was cryopreserved. Percoll-interphases were collected and rested overnight at 0.5×10e6 cells/ml in TexMACS medium (Miltenyi Biotec) plus 25 mM HEPES (pH 7.2), L-glutamine (Lonza, Köln, Germany), 50 mM beta-mercaptoethanol (ThermoFisher Scientific, Waltham, MA, USA), and 10% autologous serum. Tumor- and lung cells in the pellets were cryopreserved after resuspension. CD3-positive, CD4-positive, CD8-positive leucocyte fractions were isolated from TILs and lung-infiltrating cells using magnetic beads (Miltenyi Biotec) or FACS (BD FACS Melody, Heidelberg, Germany).

### TCR-repertoire sequencing and TIL single-cell RNA-sequencing

Genomic DNAs isolated from sorted TILs and lung-infiltrating lymphocytes were subjected to TCRSeq as described previously (Seitz 2015, Lennerz 2025)[6, 23]. Briefly, gDNAs from T-cell subpopulations (CD3, CD4, CD8,) isolated using the QIAamp blood kit (Qiagen, Hilden Germany) were used for the generation of NGS libraries by a two-step PCR protocol. Single cell cDNA-libraries were generated exclusively from CD3-positive or CD8-positive TIL single cell suspensions by use of the 10X Genomics® GemCodeTM Technology (10X Genomics B.V., Leiden, The Netherlands). Single T-cells were processed using the 10x Genomics Chromium Next GEM Single Cell V(D)J Reagent Kit in combination with the Chromium Single Cell V(D)J Enrichment Kit (Human) according to the manufacturer’s recommendations. Following clean-up, libraries were analyzed by Illumina next generation sequencing (StarSEQ GmbH, Mainz, Germany).

### Synthesis and cloning of TCR-encoding DNA-constructs

TCRα-chain and TCRβ-chain coding sequences were designed as bicistronic constructs connected by P2A-linker, ordered as G-blocks from IDT and cloned into gamma-retroviral expression vector pMX-puro as described (Lennerz 2025)[6]. A chimeric construct design was chosen with the human TRAC and TRBC domains substituted by murine Trac and Trbc sequences.

### Genetic engineering of primary T-cells

From healthy donor derived Buffy Coats, T-cells were isolated by density gradient centrifugation using Ficoll (Sigma-Aldrich, Taufkirchen, Germany) and sorting via magnetic bead separation according to the manufacturer’s protocol (Miltenyi Biotec). After activation with plate-bound (30 ng/μl) monoclonal antibody OKT-3 (anti-CD3), T-cells were subjected to CRISPR/CAS9-mediated knockout of endogenous TCRs. Ribonucleoprotein (RNP) complexes (IDT, Coralville, USA) delivered by Human T-cell Nucleofector™ Kit (Lonza, Basel, Switzerland) targeted the constant regions of both, the α- and β-TCR chains, by two specific crRNAs. The TRBC-crRNA and TRAC-crRNA were designed by the Alt-R Custom Cas9 crRNA Design Tool (IDT) as described (Lennerz 2025)[6]. In combination with the Alt-R® CRISPR-Cas9 tracrRNA (IDT) at 1:1-ratio, two different gRNA complexes were formed and subsequently combined with recombinant Cas9 (IDT) to form RNP complexes. Per knock-out reaction, 4×10e6 T-cells underwent nucleofection in Nucleofector® Solution supplemented with 1 μM Alt-R® Cas9 electroporation enhancer (IDT) and 4 μM of RNPs. For T-cell nucleofection program T-023 on a Nucleofector™ 2b Device (Lonza) was used. Thereafter, T-cells were maintained in culture at 1×10e6 cells/ml Panserin complete medium (plus 600 U/ml IL-2). Four to six days later, flow cytometry was used to assess TCR-KO efficiency.

For stable recombinant TCR knockin, γ-retroviral particles encoding the chimeric TCRs were produced for genetic engineering of primary T-cells as described (Lennerz 2025)[6]. Briefly, T-cells from healthy donor buffy coats after activation and endogenous TRAC/TRBC-KO were infected with retroviral particles encapsulating the pMX/TCR expression constructs. For production of the particles, Phoenix-ampho packaging cells were seeded at 1,3×10e6 cells per 100mm plate and after 24 hours, the Phoenix ampho cells were co-transfected with the following vectors: 5μg pCOLT-GALV, 5μg pHIT60 and 10μg pMX/TCR. Fugene-6 (Promega Madison, WI, USA) was used for transfection according to protocol. The next day, transfection medium was substituted by T-cell medium and virus-containing supernatant was harvested after 16 more hours following pelleting the cells by centrifugation. To generate TCR-T cells, activated endoTCR-KO cells were spin-inoculated with T-cell medium containing viral particles. 2×10e6 T-cells were infected per reaction and resulting TCR-T cells were expanded under selection with puromycin (1μg/ml, Sigma Aldrich).

### Culture of primary T-cells, TCR-T cells and tumor cell lines

Primary T-cells, TCR-T cells and tumor lines were grown in incubators at 37°C, 5% CO2, >85% humidity. MOR/CPR cells, NCI-H1703 cells, NCI-H1792 cells, NCI-H661 cells, NCI-H358 cells, NCI-H226 cells, and NCI-H2030 cells were purchased from ATCC or EACC and cultured under recommended conditions in the media suggested by the providers. NCI-H460 cells and MZ-LC-16 cells (received from Dr. Patrizia Haenel and Dr. Sigrid Horn, UMC, Mainz, Germany) were maintained in RPMI-1640 supplemented with 10% FBS, and 1% penicillin/streptomycin (RPMI+, Sigma Aldrich, Taufkirchen, Germany). Culture medium for primary T-cells was Panserin-413 (PAN-Biotech, Aidenbach, Germany) supplemented with 10% heat-inactivated pooled human serum (provided by the blood bank of UMC Mainz, Germany), 1% Penicillin/Streptomycin (Sigma Aldrich), and recombinant human (rh)IL-2 (Novartis, Basel, Switzerland) at concentrations ranging from 250-600 IU/mL. To check for authenticity and absence of contamination, cell lines were tested by STR-analysis and mycoplasma on a regular basis.

### Flow cytometry

T lymphocyte subpopulations were stained with the monoclonal antibodies listed in the key resource table. Expression of chimeric recombinant TCRs and differentiation from endogenous TCRs in TCR-T cells was accomplished by staining with TCR α/β (murine Trac) (Origene, cat. no. CL075F) and TCR PAN αβ (human TRAC, Beckman Coulter, Brea, USA). Antibody-labeled cells were analyzed on a FACS Canto II or Melody instrument (BD Biosciences). The resulting data (fcs-) files were exported and re-analyzed using FlowJo 10 analysis software (BD Biosciences).

### IFN-γ ELISpot assays

TCR-T cell response analyses by IFN-γ ELISpot assays were performed as shown before (Lennerz et al., PNAS 2005)[42]. Following genetic engineering, TCR-T cell populations were cultured for some weeks and aliquots cryopreserved every week. TCR-T cells were ready for application in functional assays when they showed chimeric TCR-expression ≥50%. TCR-T cells tested by ELISpot assays were taken from continuous cultures or after thawing of cryopreserved aliquots. The latter TCR-T cells were rested overnight before testing. TargeT-cells were tumor cell lines (50,000 cells/well), or autologous tumor- and normal tissue cell suspensions (20,000/well). They were co-incubated with TCR-T cells (2,000-10,000 chimeric TCR-positive cellsr as shown in γ antibody-coated Multiscreen HTS plates (Merck-Millipore, Darmstadt, Germany) overnight (16-20h). OKT3-antibody (purified from hybridoma, 400ng/ml) was co-coated in control-wells together with the anti-IFN-γ-antibody as positive reaction control. Reactions were generally set-up in replicates (duplicates or triplicates). Pan-HLA class I antibody W6/32 and anti-HLA-A*02-antibody MA2.1 (both purified from hybridoma supernatants) were used to block pMHC-specific recognition of targeT-cells by TCR-T cells. After overnight-incubation (∼20h), all cells were discarded using detergent buffers and tests developed according to protocol (Lennerz 2005)[42]. Washed and dried multiscreen-plates were scanned and analyzed by ImmunoSpot Analyzer S5 Versa with ImmunoSpot software 7.0.15.1 (CTL Europe, Bonn, Germany).

## Key Resources Table

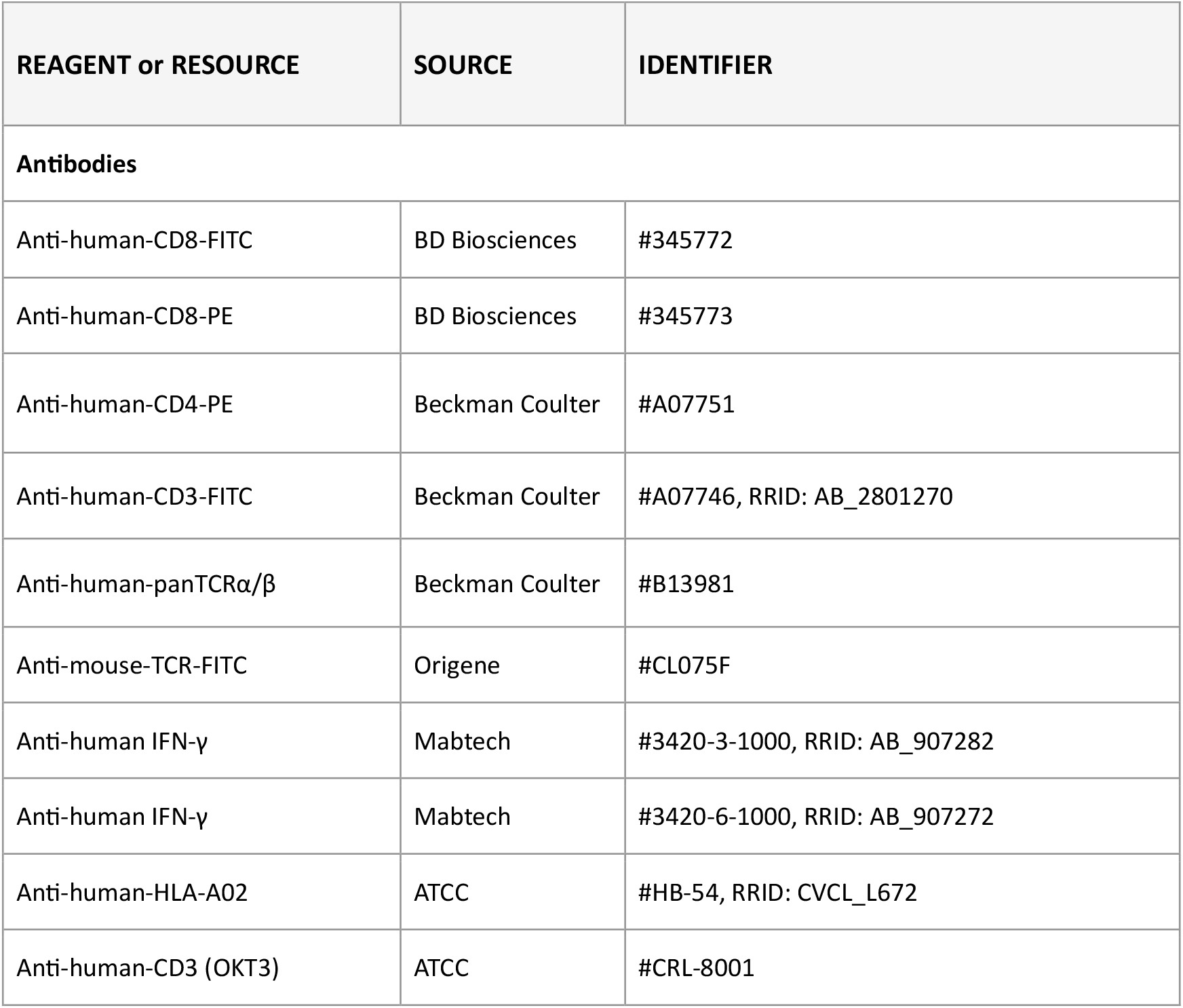
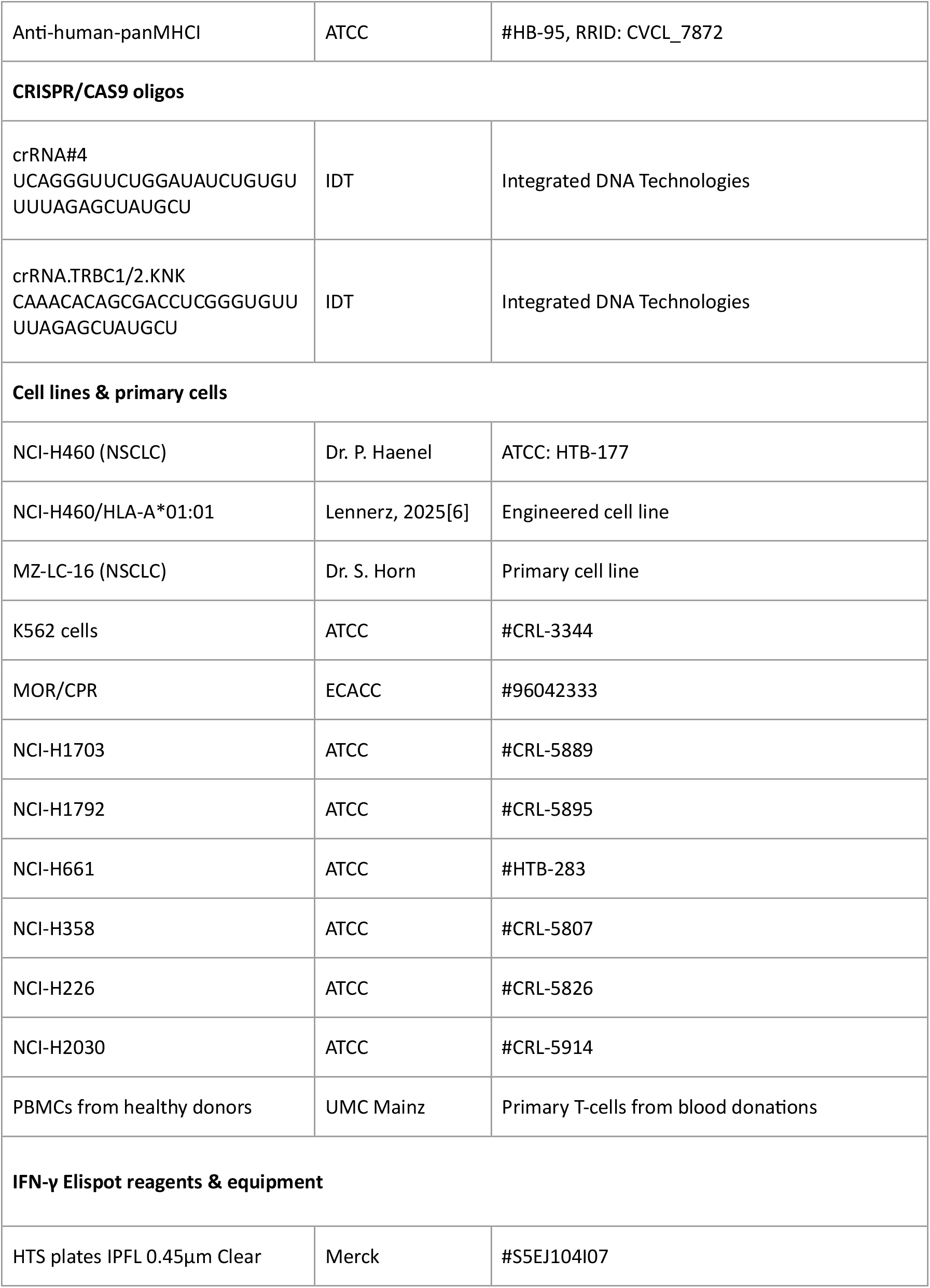
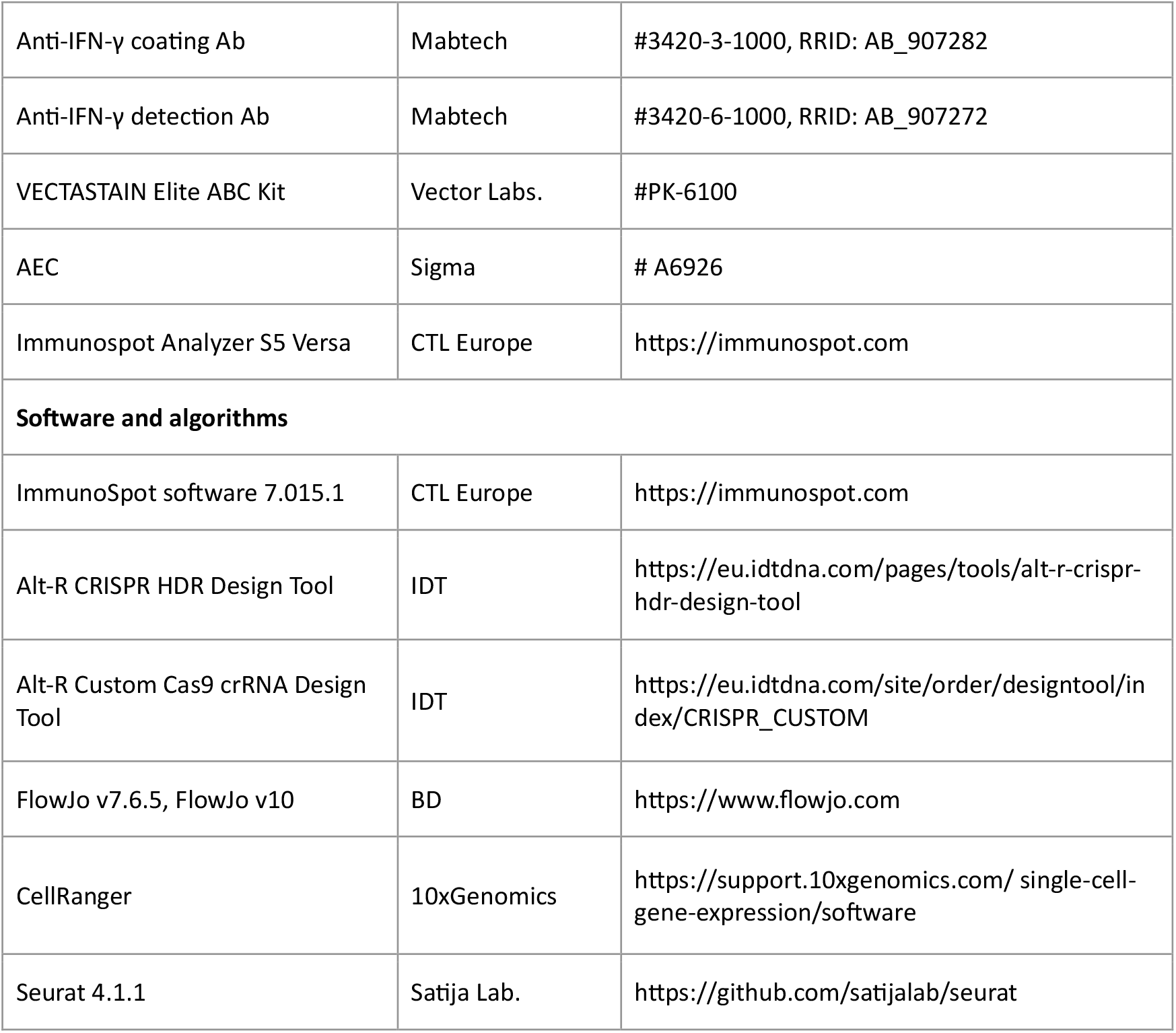

### Materials availability

This study did not generate new unique reagents.

## Data and code availability

Requests for additional data and resources should be directed to the lead contact under.

## Acknowledgments

We are grateful to Dr. Patrizia Haenel and Dr. Sigrid Horn (University Medical Clinic UMC, Mainz, Germany) for the provision of the NSCLC cell lines NCI-H460 and MZ-LC-16. We thank the blood bank of UMC Mainz for providing buffy coat preparations from whole blood of healthy donors. We are grateful to Prof. Dr. Thomas Wölfel from University Medical Center (UMC) in Mainz, Germany, for very helpful remarks and discussions concerning the manuscript. Special thanks for their cooperation in TCR analysis of NSCLC samples are due to Drs. Paul Adam, David Weismann and Samuel Lukowski from Boehringer Ingelheim RCV, GmbH, Vienna, Austria. We would like to express our gratitude to the Charité 3R and clinical teams for their assistance with tissue collection. Parts of the present work are components of CD’s doctoral thesis (Faculty 10 – Biology, Johannes Gutenberg University Mainz, Germany). We also thank the EU and the Investitionsbank Berlin (IBB) for supporting parts of the work on colorectal and breast cancer by the EFRE and ProFIT funding instruments.

## Author contributions

Conceptualization: SH, RH, KG, VL

Surgery and Pathology: SEg, CB, HH, SEl, VS, BH

Methodology: CD, AD, SS, CW, NG, SH, KG, DB, VS, BH

Investigation: KG, DB, VL, SH, HH, RH, JG

Visualization: VL, CD, KG, DB, SH

Project administration: SH, VL Supervision: VL, SH, RH

Writing – original draft: SH, RH, VL

Writing – review & editing: SH, RH, VL, VS, KG, JG

